# *In vitro* transcription using psychrophilic phage VSW-3 RNA polymerase

**DOI:** 10.1101/2020.09.14.297226

**Authors:** Heng Xia, Yixin Jiang, Rui Cheng, Bingbing Yu, Xueling Lu, Hui Wu, Bin Zhu

**Affiliations:** Key Laboratory of Molecular Biophysics, the Ministry of Education, College of Life Science and Technology and Shenzhen College, Huazhong University of Science and Technology, Wuhan, Hubei 430074, China

## Abstract

RNA research and applications were underpinned by *in vitro* transcription (IVT), while the RNA impurity resulted from the enzymatic reagents severely impede downstream applications. To improve the stability and purity of synthesized RNA we had characterized a novel single-subunit RNA polymerase (RNAP) encoded by a psychrophilic phage VSW-3 from plateau lake to produce RNA at low temperature. The VSW-3 RNAP is capable of carrying out *in vitro* RNA synthesis at low temperature (4-25°C) to reduce RNA degradation and alleviate the need of costly RNase inhibitor. Compared to routinely used T7 RNAP, VSW-3 RNAP provides comparable yield of transcripts, but is insensitive to class II transcription terminators and synthesizes RNA without redundant 3’ -cis extension. More importantly, through dot-blot detection with the J2 monoclonal antibody, we found that the RNA products synthesized by VSW-3 RNAP contain much lower amount of or virtually no double-stranded RNA (dsRNA) by-products, which are significant in most T7 RNAP products and may cause severe cellular immune response. Combining these advantages, the VSW-3 RNAP is an advantageous enzyme for IVT, especially to produce RNA for *in vivo* use.

## INTRODUCTION

DNA-dependent RNA polymerases (RNAP) specifically recognize transcriptional promoter sequences and transcribe the DNA template sequence into RNA *in vivo* and *in vitro* ^1^. However, for biotechnological use to produce large amount of RNA *in vitro*, only the single-subunit RNAPs (ssRNAPs) encoded exclusively by a group of short-tailed bacteriophages can be considered due to their simplicity and high efficiency. In the past decades, ssRNAPs from *Enterobacteria* phage T7 of T7-like viruses cluster and *Salmonella* phage SP6 of SP6-like viruses were major enzymatic reagents for *in vitro* transcription (IVT)^2, 3^. And recently T7 RNAP almost dominated IVT due to extensive research and development on it^4, 5^. However, the RNA synthesized by T7 RNAP is generally contaminated with cis-extended larger products, degraded or abortive products shorter than the DNA-encoded product^6^, and more or less but always existing dsRNA by-products resulted from non-specific transcription initiation at non-template DNA strands, the dsRNA by-product may mimic viral genome structure and is especially harmful for *in vivo* applications^7, 8^. In 2013 and 2019, we characterized representative ssRNAPs from the newly categorized P60-like marine viruses and phiKMV-like viruses, namely Syn5 and KP34 RNAP^9, 10^, they enriched the IVT toolbox by demonstrating advantages over T7 RNAP in IVT including higher processivity (Syn5)^11^, insensitivity to Class I T7 terminator (Syn5)^11^, higher product 3’ homogeneity (KP34)^10^, and higher efficiency to incorporate modified nucleotides (Syn5)^12^. However, in current conditions neither Syn5 nor KP34 RNAP can match the high RNA yield of T7 RNAP. These works encouraged us to further investigate novel ssRNAPs by leveraging the surge in viromes to identify natural variants that could foster innovations in the rapidly progressing RNA research and medicine^13–15^.

In this work, we planned to identify an ssRNAP from newly characterized cold-active bacteriophage that can efficiently synthesize RNA at low temperatures to reduce the RNA degradation caused by environmental ribonuclease contamination and prolonged incubation. We targeted the *Pseudomonas fluorescens (P. fluorescens)* bacteriophage VSW-3 of which the genome sequence information (GB: KX066068.1) was released in 2017^16^. The lytic psychrophilic bacteriophage VSW-3 was isolated together with *P. fluorescens* SW-3 cells from the Napahai located in Shangri-La County in the southwest of China^17^. In laboratory the VSW-3 phage infected *P. fluorescens* forms 2 mm clear plaques at 4°C after 48 hours^17^. The genome dsDNA of VSW-3 is 40,556 bp long and contains 46 open reading frames^17^. Together with T7, Syn5, and KP34, VSW-3 belongs to the short-tailed phage *Autographivirinae* subfamily.

We had successfully purified the VSW-3 RNAP (798 aa), identified its promoters and established its transcription system *in vitro*. As expected, VSW-3 RNAP efficiently produces RNA transcripts in the temperature range between 4-25°C, and at 25°C its maximum yield is comparable to that of T7 RNAP. Unlike all known ssRNAPs, the VSW-3 RNAP is insensitive to class II transcription terminators^18, 19^, and the RNA products are free of 3’-end extension^6, 10^. Most importantly, in contrast to T7 RNAP products, dsRNA by-products are barely detectable in VSW-3 RNAP IVT products. In addition, the Y578F mutant of VSW-3 RNAP efficiently incorporates 2’-fluoro-dATP and 2’-fluoro-dUTP into RNA^20, 21^.

## MATERIALS AND METHODS

### Materials

Oligonucleotides were synthesized by Genecreate Company, DNA purification kits from Axygen and Ni-NTA resin were from Qiagen. Preparative Superdex S200 for gel filtration was from GE Healthcare. The Gibson assembly kit, T4 RNA ligase I, Recombinant inorganic pyrophosphatase, rNTPs, DNase I, Apyrase, T7 RNA polymerase (50 U/μl), Low Range ssRNA Ladder and Monarch RNA Cleanup kit were from New England Biolabs. 2’-fluoro-dNTPs were from TriLink BioTechnologies. Reverse transcriptase kit and PrimeSTAR Max DNA Polymerase were from TaKaRa. RiboLock RNase Inhibitor and RNase A (10 mg/mL) were from Thermo Scientific. J2 monoclonal antibody (mAbs) was from English & Scientific Consulting. ImmobilonTM-Ny+ Membrane was from Millipore. X film was from Kodak. DNA marker and ECL regent were from Thermo Scientific. HPLC reversed-Phase column PLRP-S was from Agilent.

### VSW-3 RNAP expression and purification

The predicted coding sequence for VSW-3 RNAP or its Y578F mutant was inserted into pCold vector harboring N-terminal His-tag with Gibson Assembly Cloning Technology^22^. After transformation, the BL21(DE3) cells are cultured in 1 L LB medium containing 100 mg/ml ampicillin at 37°C until OD600≈0.8, incubation temperature was then reduced to 10°C and 0.2 mM IPTG was added to induce VSW-3 RNAP expression for 24 hours. The cells were harvested and resuspended in 25 mM Tris-HCl pH 7.5, 300 mM NaCl, 0.5 mM DTT, then lysed by three cycles of freeze-thaw in the presence of 0.5 mg/mL lysozyme. Supernatant was collected after centrifugation at 18000 g and 4°C for 1 hour and filtrated with 0.45 μm filter, loaded onto Ni-NTA agarose column pre-equilibrated with 10 volumes of elution buffer (25 mM Tris-HCl pH 7.5, 300 mM NaCl), then the column was washed with 10 volumes of elution buffer containing 20 mM and 50 mM imidazole, respectively, and the majority of His-tagged VSW-3 RNAP was eluted by elution buffer containing 100 mM imidazole. Collected elutes were concentrated to 2 ml with Ultra-15 Centrifugal Filter Units (Millipore) and loaded onto a 200 ml preparative Superdex S200 column for gel filtration chromatography. At last, VSW-3 RNAP was dialyzed against buffers containing 50 mM Tris-HCl pH 7.5, 100 mM NaCl, 1 mM DTT, 0.1 mM EDTA, 0.1% Triton X-100 and 50% glycerol, and then stored at −20°C. Protein concentration of VSW-3 RNAP was determined by Bradford protein quantitative kit (Bio-rad), and protein purity and concentration were analyzed along with T7 RNAP (New England Biolabs, #M0251S) by 10% SDS-PAGE gel electrophoresis stained with Coomassie blue (Bio-rad).

### VSW-3 RNAP promoter determination

In order to identify the promoter sequence and the precise transcription initiation site for VSW-3 RNAP, we first inserted the DNA fragment containing predicted promoter sequence (5’-TTAATTGGGCCACCTATAGTA-3’) into pUC19 plasmid between the *Bam*HI and *Xba*I sites with Gibson Assembly method. The plasmid was linearized by *Nde*I and purified by Axygen DNA recovery kit before serving as IVT template.

Initially we used the routine T7 RNAP IVT conditions for VSW-3 promoter determination, the 10 μl reaction contains 40 mM Tris-HCl pH 7.9, 6 mM MgCl_2_, 2 mM spermidine, 1 mM DTT, 35 ng/μl linearized pUC19 plasmid,1.5 U/μl RNase Inhibitor, 0.2 μM inorganic pyrophosphatase, 0.5 mM each of the four NTPs and 0.2 μM VSW-3 RNAP. IVT reaction was carried out at 20°C overnight. Then 1 μl DNase I was added into reaction mixture and incubation was extended for 30 min at 37°C to remove template DNA. The transcripts (pUC19-RNA) were then purified with Monarch RNA Cleanup kit.

5’-RACE of the pUC19-RNA from above step began with the RNA 5’ mono-phosphorylation treatment using the Apyrase according to New England Biolabs manual. 0.5 μg of mono-phosphorylated pUC19-RNA was ligated intermolecularly by T4 RNA ligase I in a 20 μl reaction mixture. The ligated RNA was reverse transcribed to cDNA by reverse transcriptase (TaKaRa). A pair of primers were designed: the forward primer (5’-TCGCGCGTTTCGGTGATGACGG-3’) is located at 184 nt upstream of the 3’-end of pUC19-RNA, and the reverse primer (5’-CTGATTCTGTGGATAACCGTATTAC-3’) is about 368 nt downstream of the 5’-end of pUC19-RNA. With these primers and cDNA, PCR was conducted using PrimeSTAR Max DNA Polymerase from TaKaRa. The PCR products were checked by agarose gel electrophoresis and inserted into pET28a plasmid between the *Bam*HI and *Eco*RI sites with Gibson Assembly method, then the assembled plasmids were transformed into DH5α competent cells and 5 colonies were picked for Sanger sequencing.

In order to determine the 5’ boundary of VSW-3 RNAP promoter, by annealing complementary DNA oligonucleotides, we constructed dsDNA templates with gradual truncation at the 5’ end of putative VSW-3 RNAP promoter. The constructed DNA templates each contains 18 bp, 17 bp, 16 bp, or 15 bp of the putative VSW-3 RNAP promoter upstream of the coding sequence of a 40 nt RNA product (Table S1). The 10 μl transcription reactions were carried out as described above with 4 μM annealed dsDNA templates at 20°C overnight, then 2 μl reaction product was mixed with 6 μl 2x denaturing loading buffer (95% formamide, 0.02% SDS, 1 mM EDTA, 0.02% bromophenol blue and 0.01% xylene fluoride) and 4 μl H_2_O, heated at 85°C for 2 min and immediately placed on ice for 2 min, then 10 μl sample mixture was loaded onto a 12% native TBE PAGE gel. Electrophoresis was at 100 V for 1 hour and gel was then stained with ethidium bromide before analyzed with UVsolo touch (Analytik Jena).

### IVT condition screening

A pair of primers (Table S1) were designed for PCR to prepare transcription template of cas9 RNA from a cas9-coding plasmid (addgene: 72247, T7p-cas9 plasmid), of which the T7 RNAP promoter was replaced with VSW-3 RNAP promoter (VSW-3p-cas9 plasmid). Reaction mixtures containing 0.2 μM VSW-3 RNAP, 35 ng/μl cas9 DNA template, 40 mM Tris-HCl pH 8.0, and 2 mM spermidine, 1.5 U/μl RNase Inhibitor, 0.2 μM inorganic pyrophosphatase, the concentrations of MgCl_2_ (0 mM, 1 mM, 2 mM, 3 mM, 4 mM, 5 mM, 6 mM, 7 mM, 8 mM, 9 mM, 10 mM, 12 mM, 14 mM, 16 mM, 18 mM, 20 mM) in combination with various concentrations of NTPs (0.5 mM, 1 mM, 2.5 mM, 4 mM, 5 mM) were screened for optimal RNA yield. DTT concentrations (1 mM, 5 mM, 20 mM) were also screened for a stable and high-yield transcription buffer with optimal MgCl_2_ and NTP concentrations.

With the optimal IVT condition (40 mM Tris-HCl pH 8.0, 16 mM MgCl_2_, 5 mM DTT, 2 mM spermidine, 4 mM NTPs, 35 ng/ul of cas9 DNA template, 1.5 U/μl RNase Inhibitor, and 0.2 μM inorganic pyrophosphatase), we screened enzyme concentrations (0.001 μM, 0.003 μM, 0.01 μM, 0.03 μM, 0. 1 μM, 0.15 μM and 0.3 μM) at 20°C overnight for optimal RNA yield. Finally, we screened incubation temperature (4°C, 10°C, 15°C, 20°C, 25°C, 30°C, 37°C) in combination with various incubation time in the presence of 0.15 μM VSW-3 RNAP for maximal RNA yield.

After IVT, 1 μl reaction mixture was added directly into 3 μl 2x denaturing loading buffer (95% formamide, 0.02% SDS, 1 mM EDTA, 0.02% bromophenol blue and 0.01% xylene fluoride) and 2 μl H_2_O, heated at 85°C for 2 min and immediately placed on ice for 2 min, and 5 μl of each sample was loaded onto 1.5% TAE agarose gel for electrophoresis (100 V, 30 min). The gel was stained with ethidium bromide and gel imaging was analyzed with UVsolo touch (Analytik Jena).

### Transcription termination

A group of abortive RNA products estimated to be 1500 nt-1600 nt in length were observed in T7 but not VSW-3 RNAP products during cas9 RNA synthesis. In order to find out the accurate break-off site of T7 RNA product, 3’-RACE test was conducted as following: copGFP-RNA synthesized with VSW-3 RNAP was used as adapter after mono-phosphorylation treatment with Apyrase; 0.5 μg mono-phosphorylated copGFP-RNA was ligated to the 3’-end of 0.5 μg cas9 transcripts of T7 RNAP in a 20 μl reaction mixture using T4 RNA ligase I; After reverse transcription with random primers, cDNA was PCR amplified with forward primer (5’-GTATTGCCTAAGCACAGTTTACT-3’, 1533 nt downstream to the 5‘-end of cas9-RNA), reverse primer (5’-TAGCCCATCACGTGGCTCAGCA-3’, 187 nt downstream to the 5’-end of copGFP-RNA adapter), and PrimeSTAR Max DNA Polymerase (TaKaRa); The PCR products was checked by agarose gel electrophoresis and then inserted into pET28 plasmid between the *Bam*HI and *Eco*RI sites with Gibson Assembly method. Plasmids were then transformed into DH5α competent cells and 5 colonies were picked for Sanger sequencing.

In order to confirm that the VSW-3 RNAP is not sensitive to the Class II terminators, we inserted a Class II terminator “ATCTGTT” into the copGFP RNA coding sequence located at 433 nt downstream of the 5’-end. The 10 μl IVT reaction mixture contains 40 mM Tris-HCl pH 8.0, 16 mM MgCl_2_, 5 mM DTT, 2 mM spermidine, 35 ng/μl of VSW-3p-copGFP template, 1.5 U/μl RNase Inhibitor, 0.2 μM inorganic pyrophosphatase, 4 mM each of the four NTPs and 0.15 μM VSW-3 RNAP. After IVT, the RNA products were assayed by 1.5% agarose gel electrophoresis as mentioned above.

### Synthesis of sgRNA

DNA fragments containing the coding sequence of a sgRNA (5’-<u>GGGCACGGGCAGCTTGCCGG</u>GTTTTAGAGCTAGAAATAGCAAGTTAAAATAAGGCTAGTCCGTTATCAACTTGAAAAAGTGGCACCGAGTCGGTGCTTTTTTT-3’) targeting eGFP gene under the control of a T7 or VSW-3 RNAP promoter were inserted into pUC19 plasmid between the *Bam*HI and *Xho*l sites. Then a pair of universal amplification primers (sgRNA template-F: 5’-ATCAGGCGCCATTCGCCATTCAGG-3’ and sgRNA template-R: 5’-AAAAAAAGCACCGACTCGGTGCCACT-3’) were used for PCR amplification of the sgRNA IVT template. PCR products were cleaned with DNA purification kit (Axygen). The 10 μl IVT reaction contains 40 mM Tris-HCl pH 8.0, 16 mM MgCl_2_, 5 mM DTT, 2 mM spermidine, 35 ng/ul of VSW-3p-sgRNA or T7p-sgRNA template DNA, 1.5 U/μl RNase Inhibitor, 0.2 μM inorganic pyrophosphatase, 4 mM each of the four NTPs and 0.15 μM of VSW-3 RNAP (25°C for 12 hours) or T7 RNAP (37°C for 1 hour). After IVT, the RNA products were analyzed by 12% TBE PAGE as mentioned above.

3’-RACE was carried out to verify the sgRNA product terminal homogeneity from T7 and VSW-3 RNAP IVT. Mono-phosphorylated copGFP RNA was ligated to the 3’-end of T7 and VSW-3 sgRNA transcripts with RNA ligase I. After reverse transcription, cDNA was PCR amplified with primers (3’ RACE-sgRNA-F: 5’-GCAGCTTGCCGGGTTTTAGAGCTAG-3’ and 3’ RACE-sgRNA R: 5’-TAGCCCATCACGTGGCTCAGCA-3’). The PCR products were verified by 1.5% agarose gel electrophoresis and inserted into pET28 plasmid between the *Bam*HI and *Eco*RI sites with Gibson Assembly method. 10 colonies from each group were picked for Sanger sequencing.

In addition, the sgRNA products synthesized by VSW-3 RNAP and T7 RNAP were purified with Monarch RNA Cleanup kit (New England Biolabs) and then further purified by high performance liquid chromatography (HPLC) using PLRP-S column (4000 A, 8 μM, 4.6 × 150 mm). In order to prove that the origin of the sgRNA 3’ extension is from the RNA-dependent RNA polymerase (RdRp) activity of T7 RNAP^23^, the HPLC purified VSW-3 sgRNA products were applied as the RdRp template for T7 RNAP and VSW-3 RNAP (final sgRNA concentration: 0.5 μg/μl), other components in the reaction included: 40 mM Tris-HCl pH 8.0, 16 mM MgCl_2_, 5 mM DTT, 2 mM spermidine, 4 mM each of the four NTPs, 0.2 μM inorganic pyrophosphatase, 1.5 U RNase inhibitor and 0.15 μM T7 RNAP (37°C for 1 hour) or VSW-3 RNAP (25°C for 12 hours). Reaction products were analyzed by 12% TBE PAGE as mentioned above.

### Incorporation of modified nucleotides

In order to improve resistance of RNA to RNases, 2’-fluoro (F)-dNTPs were incorporated into RNA *in vitro* and *in vivo*^20, 21^. Replacement of a tyrosine with a phenylalanine in the T7 RNAP Y639F 24, Syn5 RNAP Y564F^12^ and KP34 RNAP Y603F^10^ mutants allow synthesis of partially modified 2’-F-RNA. Through homologous sequence alignment of VSW-3 RNAP, T7 RNAP, Syn5 RNAP and KP34 RNAP with Geneious software, we identified the equivalent amino acid Y578 in VSW-3 RNAP, constructed the VSW-3 RNAP Y578F mutant and purified the mutant enzyme with the same procedure for the wild-type (WT) VSW-3 RNAP. Efficiency of the WT and Y578F VSW-3 RNAP to incorporate four 2’-F-dNTPs and other modified nucleotide substrates was tested in reactions containing 40 mM Tris-HCl pH 8.0, 16 mM MgCl_2_, 5 mM DTT, 2 mM spermidine, 1.5 U/μl RNase Inhibitor, 0.2 μM inorganic pyrophosphatase, 4 mM each of the four NTPs (with 1 of the 4 NTPs replaced by their 2’-F-dNTP, m6ATP (replacing ATP) or 5mCTP (replacing CTP) analogs, respectively), 0.15 μM of VSW-3 RNAP or VSW-3 RNAP-Y578F mutant, and 35 ng/μl of VSW-3p-sgRNA template DNA. Reactions were carried out at 25°C for 12 hours and products were analyzed by 12% TBE PAGE as mentioned above.

### Dot blot detection of dsRNA in IVT transcripts

We conducted do-blot tests with J2 monoclonal antibody to detect dsRNA in T7 and VSW-3 RNAP IVT transcripts (sox7, tdTomato, copGFP and cas9 RNA) (Table S2). Based on T7p-cas9 and VSW-3p-cas9 plasmids as mentioned above, the cas9 coding sequence of both plasmids was replaced by either the sox7, tdTomato or copGFP gene coding sequence. The transcription templates for sox7 and tdTomato RNA were plasmid DNA linearized with *BspQ*I restriction endonuclease; transcription templates for copGFP and cas9 RNA were prepared by PCR amplification with primers Trans_Template-cas9-F and Trans_Template-cas9-R (Table S1). Sox7 RNA (1388 nt), tdTomato RNA (1689 nt), copGFP RNA (928 nt) and cas9 RNA (4258 nt) were transcribed in reactions containing 40 mM Tris-HCl pH 8.0, 16 mM MgCl_2_, 5 mM DTT, 2 mM spermidine, 1.5 U/μl RNase Inhibitor, 0.2 μM inorganic pyrophosphatase, 4 mM each of the four NTPs, 35 ng/ul of template DNA, 0.15 μM of VSW-3 RNAP (25°C for 12 hours) or T7 RNAP (37°C for 1 hour). After IVT, DNA templates were removed by DNase I and transcripts were purified with Monarch RNA Cleanup kit.

RNA transcripts (200 ng) were dropped onto Immobilon™ -Ny+ Membrane (Millipore), dried, blocked with 5% non-fat dried milk in TBS-T buffer (50 mM Tris–HCl pH 7.4, 150 mM NaCl, 0.05% Tween-20), and incubated with dsRNA-specific mAb J2 (English & Scientific Consulting) for 30 min at 25°C. Membranes were washed 2 times with TBS-T buffer and reacted with HRP-conjugated donkey anti-mouse Ig (Jackson Immunology), washed 2 times and detected with ECL Western blot detection reagent (Thermo). Fluorescence signals were captured by X film (Kodak) photosensitive development. The dsRNA (0.1 ng, 0.25 ng, 0.5 ng, 1.0 ng) used as a quantitative standard was prepared by annealing two 351 nt complementary RNA strands synthesized by VSW-3 RNAP.

## RESULTS AND DISCUSSION

### Psychrophilic bacteriophage VSW-3 RNAP and its promoter

The newly discovered VSW-3 RNAP is a single-subunit RNA polymerase from Napahai lake (with an average annual temperature of 5°C) in south-western plateau of China. It has a homology of 31% to T7 RNAP and the distance tree analysis revealed that VSW-3 RNAP has different evolution distance and direction compared with T7/SP6/Syn5/KP34 RNAP (Figure 1A). As the first RNAP characterized from cold-active bacteriophage, we successfully expressed VSW-3 RNAP at 10°C. VSW-3 RNAP is smaller than T7 RNAP in size (798 aa vs. 883 aa) and the SDS page gel analysis revealed that the size of His-tagged VSW-3 RNAP is consistent with protein size prediction (92.4 kDa) (Figure 1B).

**Figure 1.**
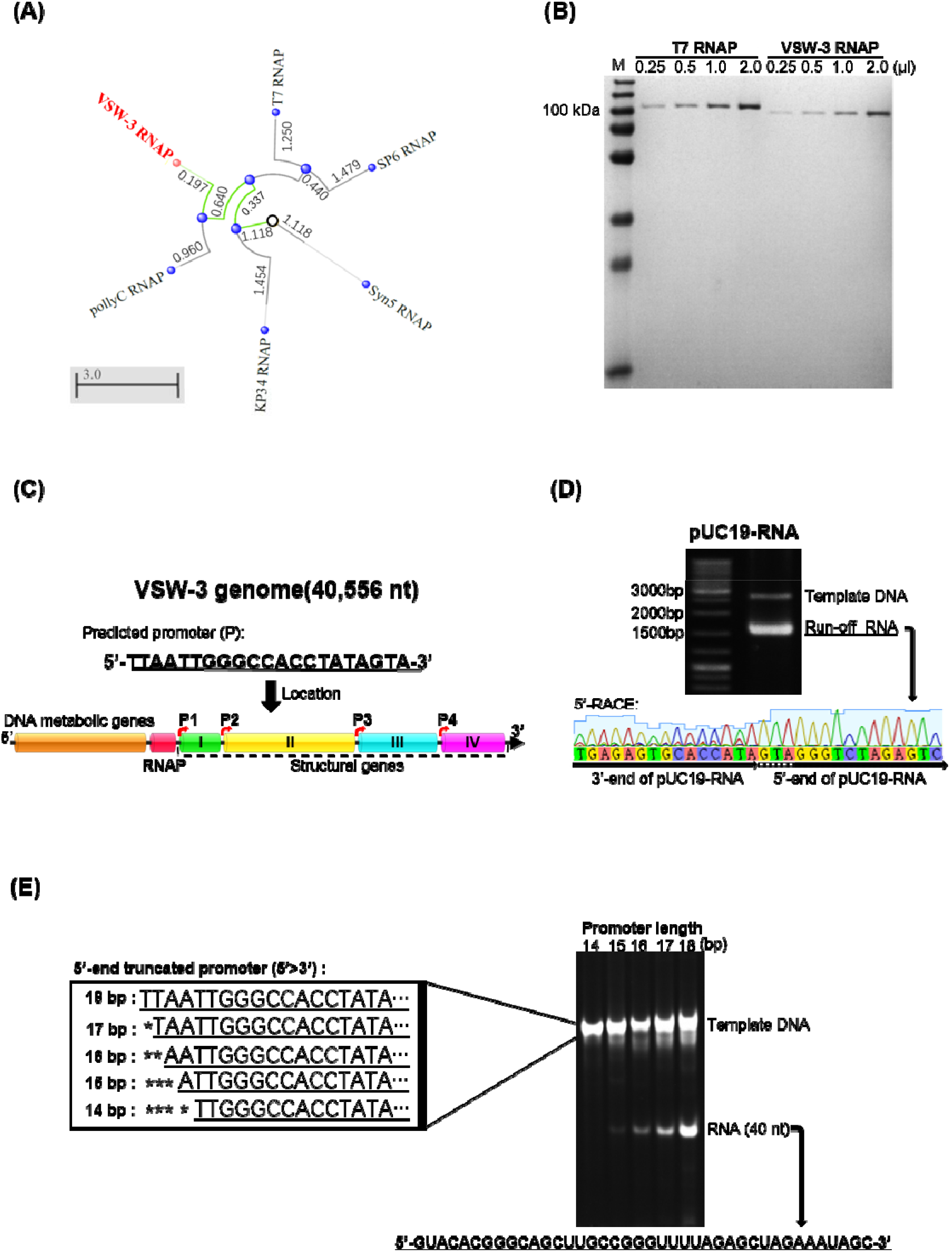
VSW-3 RNAP and its promoter. (A) Distance tree analysis of the representative ssRNAPs by Blast program. Distance from the root ‘○’: SP6 RNAP (3.374) > T7 RNAP (3.145) > KP34 RNAP (2.572) > VSW-3 RNAP (2.292) > Syn5 RNAP (1.118) suggests that VSW-3 RNAP is the second primitive after Syn5 RNAP, and evolved into a new branch of the evolutionary tree together with a predicted pollyC RNAP (3.055) from phage pollyC (YP_009622558.1). **(B)** SDS-PAGE gel analysis of purified VSW-3 RNAP (92.4 kDa including an N-terminal His-tag, 1 μM) and commercial T7 RNAP (New England Biolabs, 100 kDa, 1.5 μM), gel was stained with Coomassie blue. **(C)** Organization of phage VSW-3 genome and distribution of the predicted VSW-3 promoters (indicated by rightward arrows). (D) IVT of VSW-3 RNAP on the linearized pUC19 plasmid with an insertion of predicted VSW-3 promoter (top gel). 5’-RACE revealed that the initial nucleotides of VSW-3 RNAP transcription in the predicted promoter is “GTA” (bottom sequencing result). **(E)** IVT on 5’-truncated DNA templates (left box) to determine the accurate promoter of VSW-3 RNAP. The RNA yield with each template (right gel) suggests that the 15 bp (5’-ATTGGGCCACCTATA-3’) sequence is the minimal promoter and the 18 bp (5’-TTAATTGGGCCACCTATA-3’) sequence is the full VSW-3 promoter.

For bacteriophage T7 and SP6, transcription initiation is mainly depended on a 17 bp promoter sequence appears several times in their genomes^25–27^. However, for Syn5 and KP34^9, 10^, there are only a few promoters all over their genomes, making promoter predication difficult. Based on our previous summary of the transcription promoter distribution of representative short-tailed phages that there is at least one phage promoter between the RNAP gene and the following open reading frame^10^ and the existence of the conservative “TATA boxes”^28^, we predicted that a 21 bp nucleotide sequence (5’-TTAATTGGGCCACCTATAGTA-3’) that appears four times in VSW-3 genome most likely harbors the VSW-3 promoter. One of the predicted promoter regions follows the VSW-3 RNAP gene (13579-13599) and the other three located at 17122-17142, 28006-28026, and 34846-34866, respectively, all in the intergenic regions (Figure 1C).

Using linearized pUC19 plasmid with predicted VSW-3 promoter inserted as transcription template, the IVT activity of VSW-3 RNAP was firstly demonstrated (Figure 1D). We analyzed the IVT transcripts to confirm the VSW-3 RNAP transcriptional initiation site, 5’-RACE revealed that the transcripts start with “GTA” (Figure 1D). Gradually truncation on the 5’-end of the 21 bp putative VSW-3 promoter region revealed that the 15 bp sequence (5’-ATTGGGCCACCTATA-3’) is the minimum promoter and the 18 bp sequence (5’-TTAATTGGGCCACCTATA-3’) is the full promoter for VSW-3 RNAP (Figure 1E). The main difference between T7 (5’-TAATACGACTCACTATA-3’) and VSW-3 promoters is the middle 8 bp sequence (T7: ACGACTCA, VSW-3: TGGGCCAC).

### Optimized VSW-3 RNAP IVT system

We found that the IVT yield of VSW-3 RNAP is significantly affected by the concentration of Mg^2+^ and DTT, higher concentration of NTPs requires a corresponding higher Mg^2+^ in the transcription buffer (Figure 2A). Although the buffer contained 9 mM Mg^2+^ and 1 mM DTT could support an optimal yield of cas9 RNA in the presence of 4 mM NTPs (Figure 2A), it was highly unstable and the yield of cas9 RNA dropped significantly with this buffer stored at −20 °C for a few days (Figure 2A). Higher concentrations of DTT (5 mM and 20 mM) increases the buffer stability and meanwhile requires higher Mg^2+^ concentration to reach maximum yield of RNA (Figure 2B). Finally, we have established an efficient and stable (more than 6 months) IVT buffer for VSW-3 RNAP (40 mM Tris-HCl pH 8.0, 16 mM MgCl_2_, 5 mM DTT and 2 mM spermidine).

**Figure 2.**
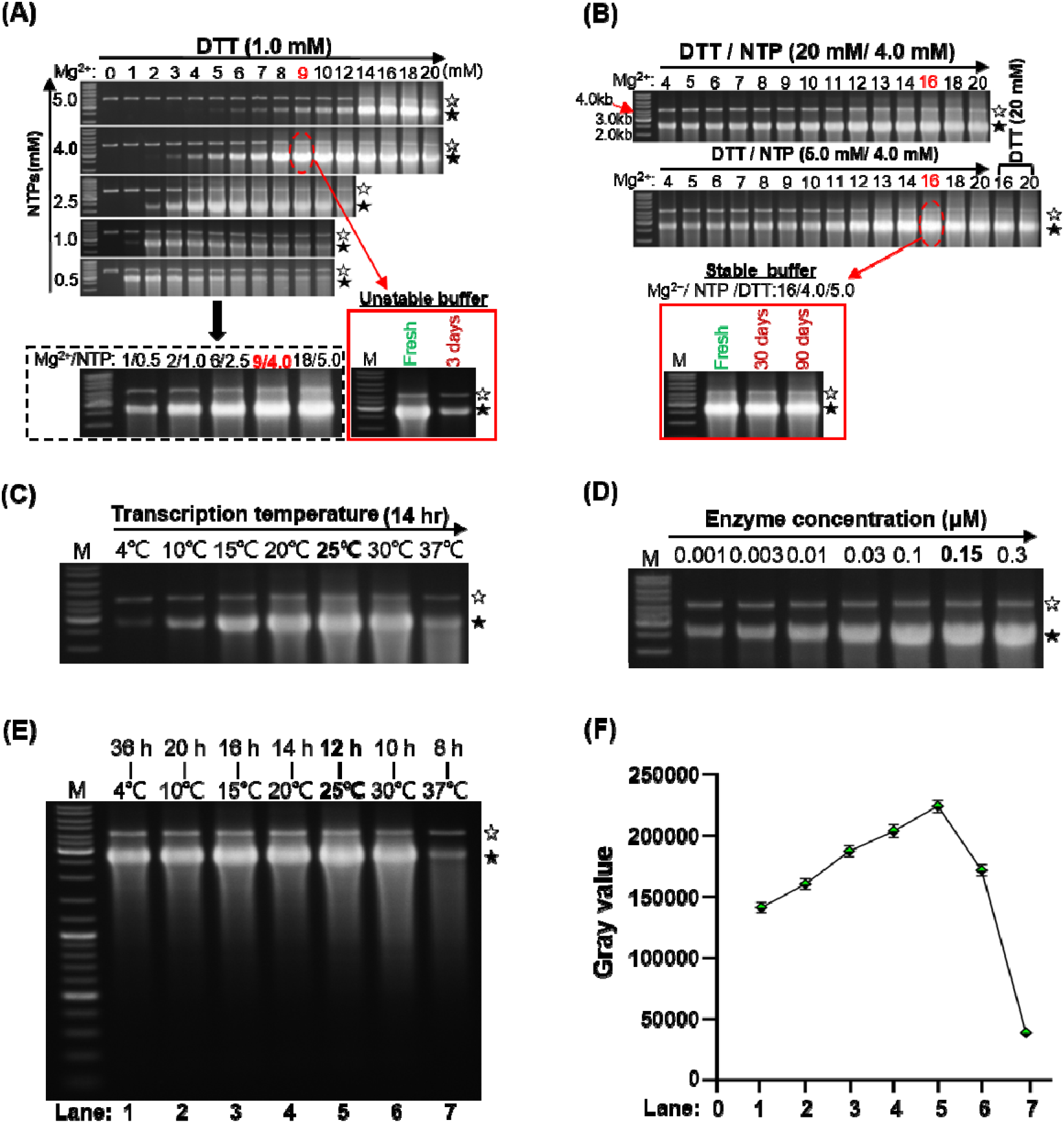
Optimal VSW-3 RNAP IVT conditions. **(A)** Screening for the optimal Mg^2+^/NTP concentration in the presence of 1 mM DTT. RNA yield with various optimal Mg^2+^/NTP concentration combination was further compared (gel in the dotted box). The stability of the optimal VSW-3 RNAP IVT buffer with 1 mM DTT was examined (gel in the solid box). **(B)** Screening for the optimal DTT/ Mg^2+^ concentration for the stable and high-yield VSW-3 RNAP IVT buffer. The stability of the high-yield VSW-3 RNAP IVT buffer containing 16 mM Mg^2+^, 4 mM NTP and 5 mM DTT was examined (gel in the solid box). **(C)** The optimal reaction temperature of VSW-3 RNAP (25°C) for maximum run-off RNA yield. **(D)** The optimal enzyme concentration of VSW-3 RNAP (0.15 μM) for maximum run-off RNA yield. **(E)** Optimal IVT yield of VSW-3 RNAP with various reaction temperature/incubation time combinations. The maximum run-off RNA yield was obtained at 25°C for 12 hours. **(F)** Gray-scale quantitation of the run-off RNA transcripts in gel **(E)**. Diagram was made using GraphPad Prism. In all gels the bands corresponding to DNA templates were indicated by empty stars and the bands corresponding to run-off RNA transcripts were indicated by filled stars.

VSW-3 RNAP produces RNA within a wide range of temperature (4-37°C) *in vitro*, with the highest yield at 25°C (Figure 2C). While at lower temperatures, the yield increases as incubation time extended (Figure S1). The optimal enzyme concentration of VSW3 RNAP is 0.15 μM (Figure 2D), similar as that for T7 RNAP. At 37°C, the optimal temperature for T7 RNAP, the yield of VSW-3 RNAP is even lower than that at 4°C (Figure 2E). Maximum IVT yield of RNA for VSW-3 RNAP was obtained at 25°C after a 12-hr incubation (Figure 2E and 2F). A working stock of 1.5 μM VSW-3 RNAP is stable for more than half a year at −20°C.

The optimized transcription buffer for VSW-3 RNAP (16 mM MgCl_2_, 5 mM DTT) is also optimal for T7 RNAP to achieve high yield in the presence of 4 mM NTPs, compared to the routine buffer (6 mM MgCl_2_, 1 mM DTT, New England Biolabs) (Figure S2A and S2B). At this condition the T7 RNAP reaches its maximum yield in 1 hour, and further extended incubation does not increase the yield, and even decreases the amount of run-off products (Figure S2A and S2B).

With the same optimal buffer, the same concentrations of enzyme and substrates, the maximum yield of VSW-3 (25 °C, 12 hours) and T7 RNAP (37 °C, 1 hour) is comparable, while for the synthesis of long RNA such as cas9 RNA (Figure 3A), tdTomato (GenBank: KT878736.1) RNA and complete S protein RNA of SARS-CoV-2 (GenBank: NC_045512.2), the optimal yield of VSW-3 RNAP (quantified as about 70 μg per 20 μl IVT reaction for these RNA) is higher than that of T7 RNAP (quantified as about 60 μg per 20 μl IVT reaction for these RNA). We also compared the pUC19-RNA yield of VSW-3 and T7 RNAP IVT at 25 °C (thus all IVT conditions are the same for each enzyme) and consistently found that the maximum yield of VSW-3 RNAP IVT was higher than that of T7 RNAP IVT (Figure S2C).

**Figure 3.**
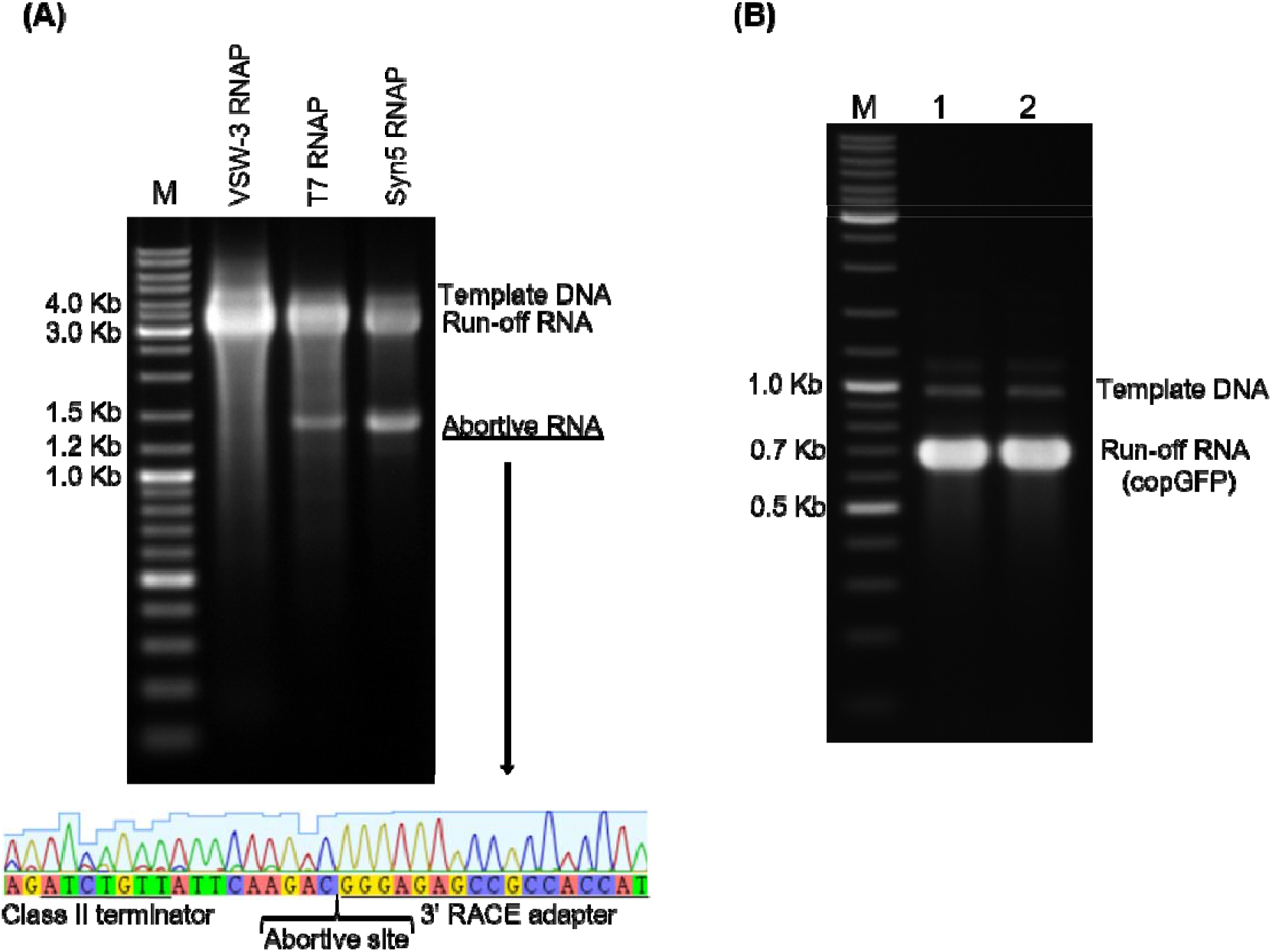
Response of ssRNAPs to Class II terminator. **(A)** Using PCR-amplified templates for cas9-RNA IVT, obvious abortive RNA transcripts were observed for T7 RNAP and Syn5 RNAP but not VSW-3 RNAP (top gel). 3’-RACE revealed that the T7 RNAP transcription was terminated 9 nt downstream of a Class II terminator “ATCTGTT” (bottom sequencing result). **(B)** VSW-3 RNAP IVT was not terminated (no additional bands comparing lane 2 with lane 1) when a Class II terminator “ATCTGTT” was inserted into the middle of the copGFP RNA coding sequence.

As we expected, IVT at lower temperature can reduce the RNA degradation by RNase. Even at 25°C for 12 hours, the degradation of VSW-3 transcripts by RNase A is significantly lower than that of T7 transcripts at 37°C for 1 hour during IVT (Figure S3). To alleviate the need of costly RNase inhibitor and strict RNase-free IVT environment, IVT with VSW-3 RNAP can be carried out at lower temperatures (such as in refrigerators) with longer incubation time (Figure 2E). Without adding RNase inhibitor into VSW-3 RNAP IVT reactions while keeping the reactions in 4-25 °C, we observed no transcripts degradation.

### Transcription termination

There are two classes (class I and class II) of transcription terminators previously known to partially stop the transcriptional elongation of T7 RNAP, the class I terminator forms a stem-loop structure and the class II terminator typically contains a 7 nt sequence (5’-ATCTGTT-3’)^18, 29, 30^. During the transcriptional assays with PCR product as transcription template for cas9 RNA (Addgene: 72247), we found that there was a group of abortive RNA products synthesized by T7 and Syn5 RNAP. According to the markers in the agarose gel, the abortive RNA products were estimated to be 1500 nt to 1600 nt in length (Figure 3A gel). Through 3’-RACE, we found that the abortive site of T7 RNAP was located 9 nt downstream of a typical class II terminator “ATCTGTT” in cas9 RNA (Figure 3A bottom). However, in the same RNA this terminator showed no effect on VSW-3 RNAP (Figure 3A gel), and a class II terminator 5’-ATCTGTT-3’ inserted into the copGFP gene didn’t cause any abortive-product synthesis by VSW-3 RNAP (Figure 3B). To our knowledge, VSW-3 RNAP is the only known ssRNAP that is naturally insensitive to the class II terminators, making it suitable to synthesize RNAs containing class II terminator to avoid abortive products.

### 3’ termini of RNA products

T7 RNAP retains the RNA-dependent RNA polymerase (RdRp) activity^23^ and catalyzes the self-templated extension on the RNA 3’ hairpin structure^6, 31^. The recently widely used single-guide RNA (sgRNA, 103 nt) is a typical RNA with terminal secondary structure. The first 20 nt sequence of the sgRNA tested in this work is a crRNA (5’-GGGCACGGGCAGCTTGCCGG-3’) targeting eGFP, the rest 83 nt is the gRNA backbone, and the secondary structure of this sgRNA was predicted with RNAfold^32^ (Figure 4A). With native 12% TBE PAGE assay, products with extended 3’ termini were observed for T7 RNAP but not VSW-3 RNAP, although the latter produced more abortive products (Figure 4B). 3’-RACE revealed that among the 10 colonies picked for T7 RNAP sgRNA products (Figure 4C), only one sequencing result matches the full-length sgRNA (103 nt), 7 sequencing results revealed there were 3’-end extra extensions up to 16 nt. The mechanism of such extension by the RdRp activity of T7 RNAP is presented in Figure 4D. The parallel sequencing results for VSW-3 RNAP sgRNA products showed no 3’ extension at all, consistent with gel analysis (Figure 4B). Among 10 sequences, 3 match the exact sgRNA sequence (103 nt), 3 miss one nucleotide (102 nt), and the other 4 showed further truncation (101 nt, 99 nt, 40 nt and 33 nt, respectively) of which the underlying mechanism is not clear (Figure 4C).

**Figure 4.**
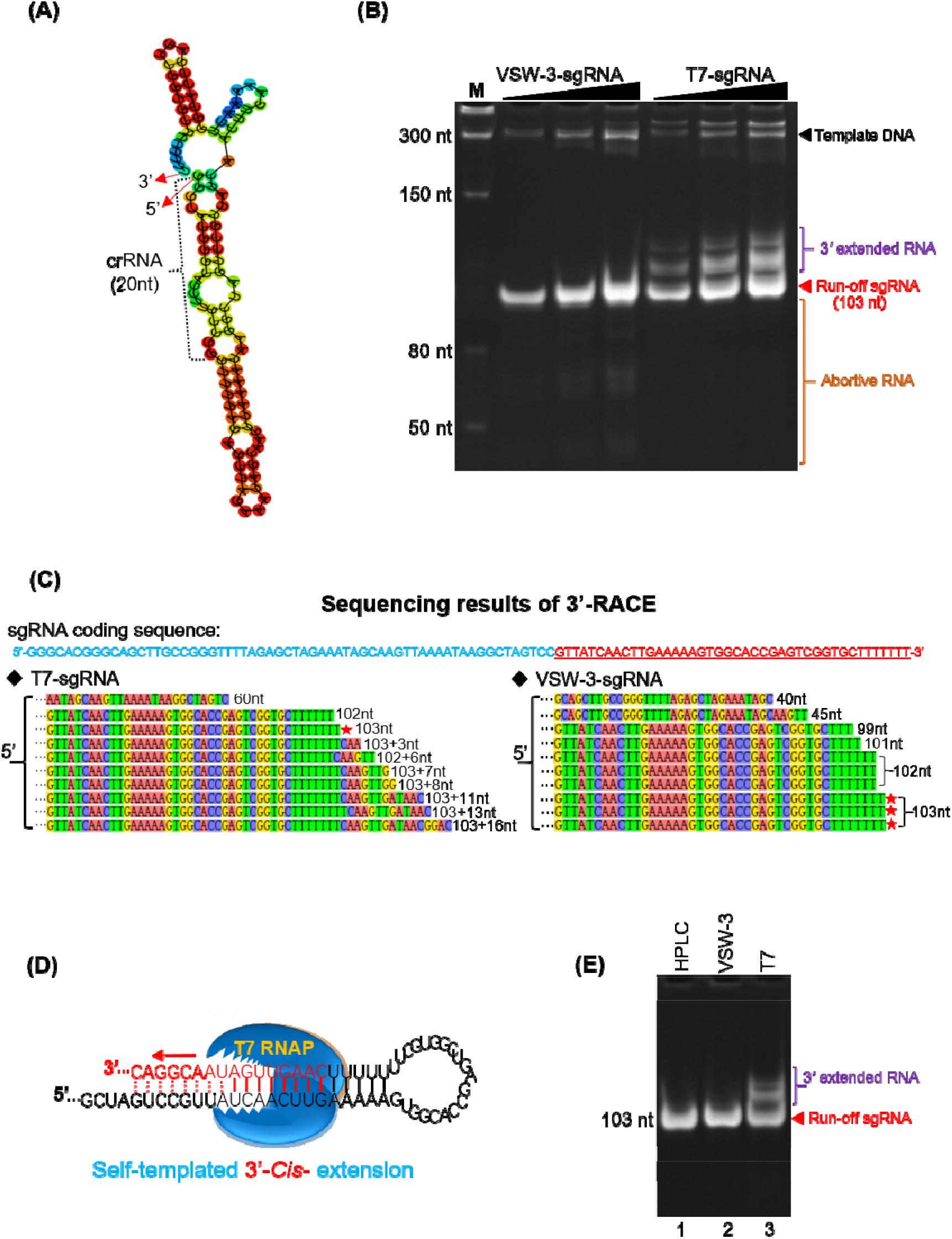
RNA 3’ extension and RdRp activity of T7 and VSW-3 RNAP. **(A)** The secondary structure of a sgRNA predicted with RNAfold software. **(B)** IVT synthesis of a sgRNA (targeting eGFP) by VSW-3 and T7 RNAP. **(C)** 3’-RACE of the sgRNAs transcripts from T7 and VSW-3 RNAP IVT. Only the 3’ region (red sequence on the top) of the full sgRNA in sequencing results was shown. The length of each sequence was noted. The sequences matching the exact run-off sgRNA (103 nt) was indicated by red stars. **(D)** Schematic showing the mechanism and origin (3’ self-templated extension by the RdRp activity of T7 RNAP) of the 16 nt 3’-extension in T7 RNAP products as in **(C)**. **(E)** T7 but not VSW-3 RNAP retains the RdRp activity to extend purified sgRNA (with terminal primer/template structure).

HPLC was applied to the sgRNA products from T7 and VSW-3 RNAP IVT. Consistent with PAGE and 3’-RACE results, HPLC chromatogram showed a relatively small peak (3’ extended sgRNA) following the main peak (run-off sgRNA transcripts) for T7 RNAP products (Figure S4A). This chromatogram peak corresponding to the 3’ extended sgRNA was not shown for VSW-3 RNAP products, instead other peaks corresponding to abortive RNA products was observed (Figure S4B).

With the HPLC-purified sgRNA product of VSW-3 RNAP as template, we examined the RdRp activity of T7 and VSW-3 RNAP (Figure 4E). Such activity was not detected for VSW-3 RNAP as the RNA template with terminal primer/template-like structure was not extended. However, T7 RNAP efficiently extended such RNA template with its RdRp activity^23^ (Figure 4E), confirming the origin of the 3’ extended transcripts in T7 RNAP IVT.

### VSW-3 RNAP mutant for incorporation of modified nucleotides

ssRNAPs usually prefer NTPs against dNTPs and 2’-F-dNTPs, however, replacement of a tyrosine with a phenylalanine in the ‘O-helix’ region such as those in the T7 Y639F^24, 33^, Syn5 Y564F ^12^, and KP34 Y603F (10) RNAP mutants weakens this discrimination and allows modified 2’-F-RNA synthesis. RNA containing ribose modifications such as 2’-F is more resistant to RNase A, which recognizes the 2’-OH of pyrimidines for cleavage, resulting in longer survival time of 2’-F-RNA *in vitro* and *in vivo*^20, 21, 34, 35^. We investigated the effect of such mutation in the VSW-3 RNAP. Homologous sequence alignment revealed the equivalent amino acid Y578 in VSW-3 RNAP (Figure 5A). The Y578F mutation decreased the yield of native RNA (Figure 5B). When UTP in the sgRNA IVT reaction was replaced by 2’-F-dUTP, the WT VSW-3 RNAP produced small amount of run-off products along with large number of abortive products, indicating that the 2’-F-dUTP decreased the processivity of VSW-3 RNAP (Figure 5C). This effect was removed by the Y578F mutation as the mutant efficiently produced run-off sgRNA with 2’-F-dUTP (Figure 5C). Unexpectedly, the incorporation of 2’-F-dCTP was only slightly improved by the VSW-3 Y578F mutation. Instead, the incorporation of 2’-F-dATP was significantly increased by the VSW-3 Y578F mutation (Figure 5C), unlike cases for KP34 RNAP Y603F and Syn5 RNAP Y564F mutants^10, 12^. Neither the WT nor Y578F VSW-3 RNAP incorporated 2’-F-dGTP. We also examined the incorporation of other modified nucleotide substrates including N6-methyl-ATP (m6ATP) and 5-methyl-CTP (5mCTP), both the WT and Y578F VSW-3 RNAP efficiently incorporated m6ATP and inefficiently utilized 5mCTP (Figure 5C).

**Figure 5.**
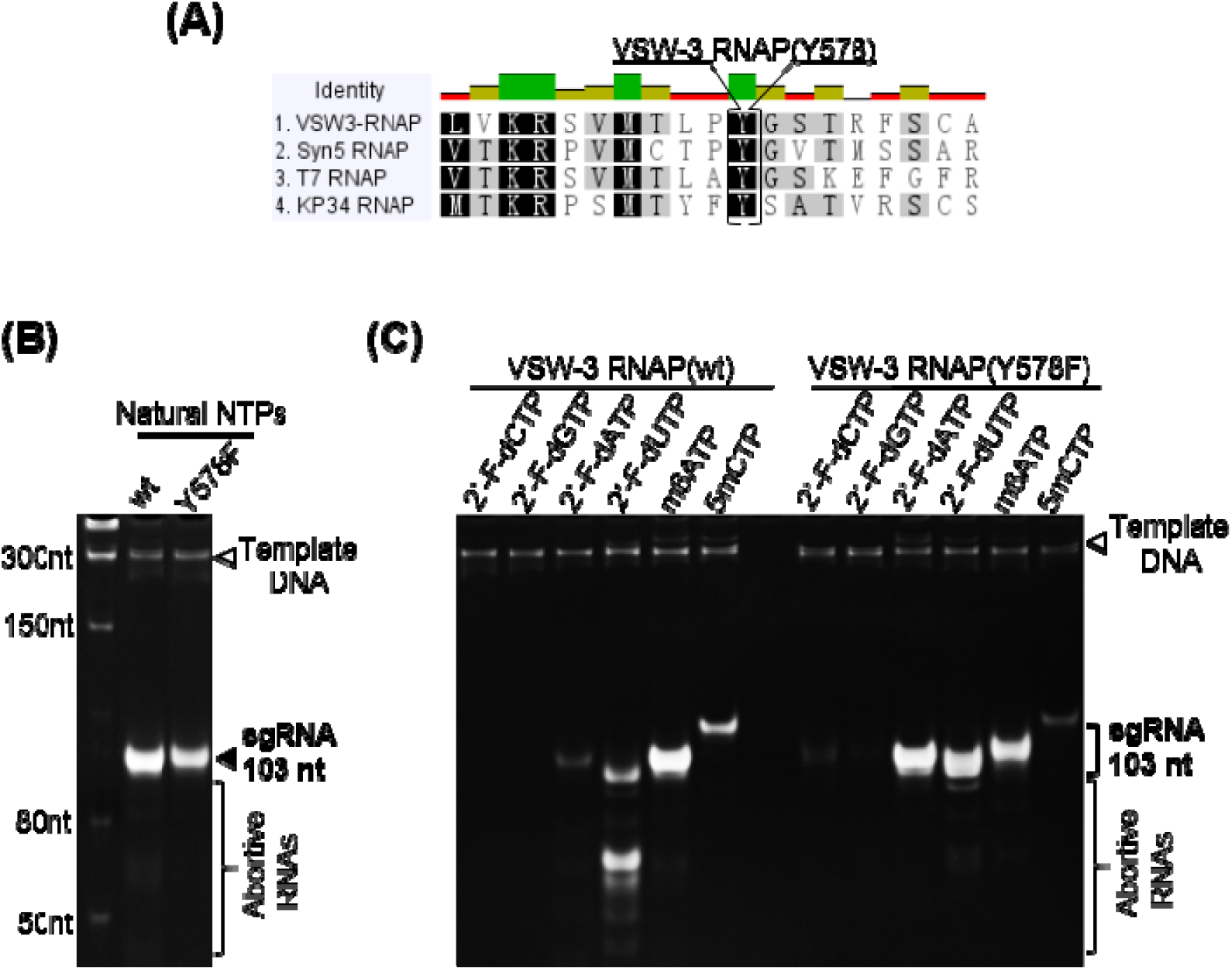
Incorporation of modified nucleotides by WT and Y578F VSW-3 RNAP. **(A)** Homologous sequence alignment of VSW-3 RNAP, T7 RNAP, Syn5 RNAP and KP34 RNAP using Geneious software identified the tyrosine (Y578) in the ‘O-helix’ region of VSW-3 RNAP. **(B)** IVT yield of a sgRNA by WT and Y578F VSW-3 RNAP was compared. **(C)** In the IVT system for a sgRNA synthesis, the incorporation of modified nucleotides by WT and Y578F VSW-3 RNAP was compared. In each reaction, one of the four NTPs was replaced by its modified analog (as indicated on top of the gel). The efficiency of incorporation was judged based on the yield of the run-off sgRNA.

### dsRNA by-products

dsRNA is another routine by-products from T7 RNAP IVT^7, 8^, resulted from annealing of plus strand RNA transcripts with minus strand non-specific RNA transcripts (T7 RNAP non-specifically recognized sequences in the DNA template that resemble T7 promoter. If the non-specific transcription initiation occurs at non-template strand, the minus strand RNA transcripts are generated)^7^. Depending on their sequences, sizes and structures, it is difficult to remove dsRNA completely through purification, even by HPLC, especially from scaled-up IVT products^5^. And the remaining dsRNA, although in small amount, may cause severe cellular immune response (by mimicking RNA virus genome component) if the RNA transcripts are applied *in vivo*^7, 8^. In some T7 RNAP IVT products, the suspected dsRNA can be observed on agarose gel as slower moving bands larger than the major run-off RNA transcripts, for example, in the case of copGFP (GenBank: KX757255.1, Table S2) RNA synthesis (Figure 6B). However, for the same RNA, the band corresponding to dsRNA was not observed for VSW-3 RNAP IVT (Figure 6B). For other transcripts, such as sox7 (GenBank: NM_031439.4, Table S2), tdTomato (GenBank: KT878736.1, Table S2), and cas9 (Addgene: 72247, Table S2) RNA, although the dsRNA from T7 RNAP IVT is not obvious on gel (Figure 6B), the dot blot assay with J2 monoclonal antibodies (mAbs) confirmed that for most T7 RNAP transcripts (except for the sox7 RNA), every 200 ng contains more than 2.0 ng dsRNA according the dsRNA quantitative standard (Figure 6C and 6D). While for VSW-3 RNAP, most examined transcripts contain barely detectable dsRNA (Figure 6C and 6D).

**Figure 6.**
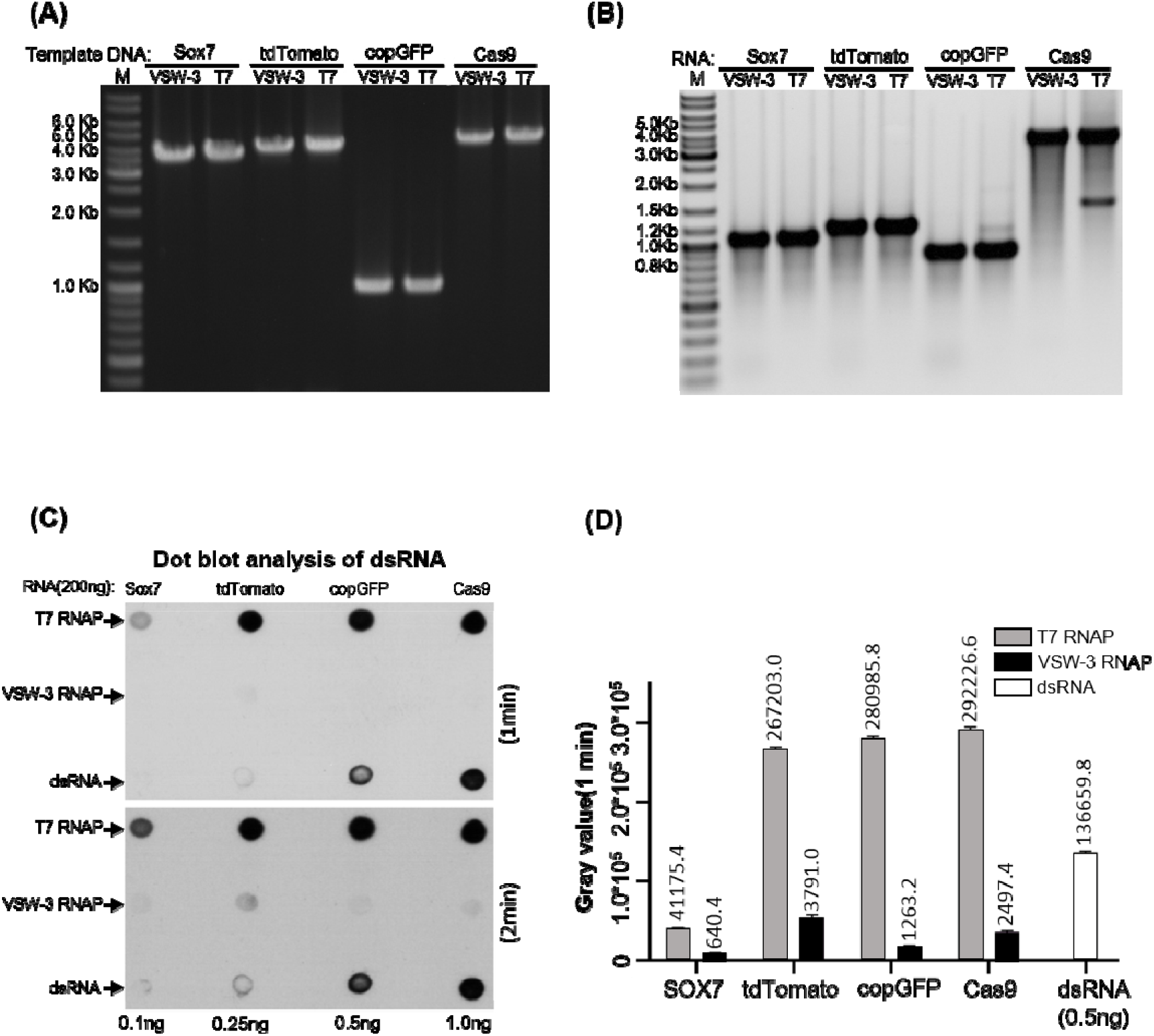
dsRNA by-products from T7 and VSW-3 RNAP IVT. **(A)** DNA templates for the IVT synthesis of various RNA as indicated on top of the gel. For each RNA, there are two DNA templates differ only in the promoter region to serve for VSW-3 and T7 RNAP IVT, respectively. DNA concentration and purity were compared in 1.5% agarose gel stained with ethidium bromide. **(B)** After template DNA was removed by DNase I treatment and purified with Monarch RNA Cleanup kit, 1μg of sox7, tdTomato, copGFP and Cas9 RNA transcribed by VSW-3 RNAP and T7 RNAP were analyzed in 1.5% agarose gel stained with ethidium bromide. The white and black colors for bands and background were converted in this gel picture to make the weak double-stranded and abortive RNA bands clearer. **(C)** Dot blot analysis of the RNA products (each 200 ng) as in **(B)** by VSW-3 RNAP and T7 RNAP with J2 monoclonal antibody. A prepared dsRNA (351 bp) was applied as quantitative standard (0.1 ng, 0.25 ng, 0.5 ng, 1.0 ng). **(D)** The gray value measurement and calculation of the X film image (top image in **(C)**) by Image J software demonstrating the level of dsRNA contamination in T7 and VSW-3 RNAP transcripts.

The VSW-3 RNAP is the first characterized ssRNAP from psychrophilic phage. With comparable RNA yield to the routinely used T7 RNAP, its ability to perform RNA synthesis at low temperature can significantly reduce the RNA degradation during prolonged synthesis and in the absence of RNase inhibitors. In combination with its other advantages including insensitivity to Class II T7 transcription terminators, higher product 3’-terminal homogeneity, and importantly, minimal dsRNA production, the VSW-3 RNAP is especially advantageous in the synthesis of IVT transcripts for *in vivo* use, such as mRNA medicine. Indeed, colleagues had communicated unprecedented cellular expression level of mRNA (without modification and HPLC purification) produced by VSW-3 RNAP in *in vivo* applications (In preparation).

## Supporting information

VSW-3 RNA polymerase-Supplementary Materials

## SUPPLEMENTARY DATA

Supplementary Data are available at NAR online.

## ACKNOWLEDGEMENT

We thank Dr. Hao Yin and Guoquan Wang of Wuhan University for helpful discussions on *in vivo* mRNA expression.

