## Supplementary material for "*In vitro* transcription using psychrophilic phage VSW-3 RNA polymerase": VSW-3 RNA polymerase-Supplementary Materials

**Table S1. DNA oligos for the IVT templates preparation.**

| **Primers for PCR amplification of the cas9 RNA IVT template** | |
| --- | --- |
| **Trans_Template-cas9-F：** | AGCTGGTTTAGTGAACCGTCAGATC |
| **Trans_Template-cas9-R：** | ACTCAATGGTGATGGTGATGATGACC |
| **DNA oligos for the construction of truncated VSW-3 RNAP promoter** | |
| **VSW3-promoter Test (18)-F:** | **TTAATTGGGCCACCTATA**GTACACGGGCAGCTTGCCGGGTTTTAGAGCTAGAAATAGC |
| **VSW3-promoter Test (18)-R:** | GCTATTTCTAGCTCTAAAACCCGGCAAGCTGCCCGTGTAC**TATAGGTGGCCCAATTAA** |
| **VSW3-promoter Test (17)-F:** | **TAATTGGGCCACCTATA**GTACACGGGCAGCTTGCCGGGTTTTAGAGCTAGAAATAGC |
| **VSW3-promoter Test (17)-R:** | GCTATTTCTAGCTCTAAAACCCGGCAAGCTGCCCGTGTAC**TATAGGTGGCCCAATTA** |
| **VSW3-promoter Test (16)-F:** | **AATTGGGCCACCTATA**GTACACGGGCAGCTTGCCGGGTTTTAGAGCTAGAAATAGC |
| **VSW3-promoter Test (16)-R:** | GCTATTTCTAGCTCTAAAACCCGGCAAGCTGCCCGTGTAC**TATAGGTGGCCCAATT** |
| **VSW3-promoter Test (15)-F:** | **ATTGGGCCACCTATA**GTACACGGGCAGCTTGCCGGGTTTTAGAGCTAGAAATAGC |
| **VSW3-promoter Test (15)-R:** | GCTATTTCTAGCTCTAAAACCCGGCAAGCTGCCCGTGTAC**TATAGGTGGCCCAAT** |
| **VSW3-promoter Test (14)-F:** | **TTGGGCCACCTATA**GTACACGGGCAGCTTGCCGGGTTTTAGAGCTAGAAATAGC |
| **VSW3-promoter Test (14)-R:** | GCTATTTCTAGCTCTAAAACCCGGCAAGCTGCCCGTGTAC**TATAGGTGGCCCAA** |

**Table S2. Sequences (5’-3’) of sox7, tdTomato, copGFP and cas9 RNA.**

sox7 RNA sequence (GenBank: NM_031439.4)

GGGAGACCCUCGAGGACAGAUCGCCUGGAGACGGCAAGAGCCGCCACCAUGAAAAGGCCGGCGGCCACGAAAAAGGCCGGCCAGGCAAAAAAGAAAAAGGGUUCUGGAGCUUCGCUGCUGGGAGCCUACCCUUGGCCCGAGGGUCUCGAGUGCCCGGCCCUGGACGCCGAGCUGUCGGAUGGACAAUCGCCGCCGGCCGUCCCCCGGCCCCCGGGGGACAAGGGCUCCGAGAGCCGUAUCCGGCGGCCCAUGAACGCCUUCAUGGUUUGGGCCAAGGACGAGAGGAAACGGCUGGCAGUGCAGAACCCGGACCUGCACAACGCCGAGCUCAGCAAGAUGCUGGGAAAGUCGUGGAAGGCGCUGACGCUGUCCCAGAAGAGGCCGUACGUGGACGAGGCGGAGCGGCUGCGCCUGCAGCACAUGCAGGACUACCCCAACUACAAGUACCGGCCGCGCAGGAAGAAGCAGGCCAAGCGGCUGUGCAAGCGCGUGGACCCGGGCUUCCUUCUGAGCUCCCUCUCCCGGGACCAGAACGCCCUGCCGGAGAAGAGAAGCGGCAGCCGGGGGGCGCUGGGGGAGAAGGAGGACAGGGGUGAGUACUCCCCCGGCACUGCCCUGCCCAGCCUCCGGGGCUGCUACCACGAGGGGCCGGCUGGUGGUGGCGGCGGCGGCACCCCGAGCAGUGUGGACACGUACCCGUACGGGCUGCCCACACCUCCUGAAAUGUCUCCCCUGGACGUGCUGGAGCCGGAGCAGACCUUCUUCUCCUCCCCCUGCCAGGAGGAGCAUGGCCAUCCCCGCCGCAUCCCCCACCUGCCAGGGCACCCGUACUCACCGGAGUACGCCCCAAGCCCUCUCCACUGUAGCCACCCCCUGGGCUCCCUGGCCCUUGGCCAGUCCCCCGGCGUCUCCAUGAUGUCCCCUGUACCCGGCUGUCCCCCAUCUCCUGCCUAUUACUCCCCGGCCACCUACCACCCACUCCACUCCAACCUCCAAGCCCACCUGGGCCAGCUUUCCCCGCCUCCUGAGCACCCUGGCUUCGACGCCCUGGAUCAACUGAGCCAGGUGGAACUCCUGGGGGACAUGGAUCGCAAUGAAUUCGACCAGUAUUUGAACACUCCUGGCCACCCAGACUCCGCCACAGGGGCCAUGGCCCUCAGUGGGCAUGUUCCGGUCUCCCAGGUGACACCAACGGGUCCCACAGAGACCAGCCUCAUCUCCGUCCUGGCUGAUGCCACGGCCACGUACUACAACAGCUACAGUGUGUCAGGAUCCCCCAAGAAGAAGAGGAAAGUCUCGAGCGACUACAAAGACCAUGACGGUGAUUAUAAAGAUCAUGACAUCGAUUACAAGGAUGACGAUGACAAGGCUGCAGGAUGAA

tdTomato RNA sequence (GenBank: KT878736.1)

GGGAGACCCUCGAGGACAGAUCGCCUGGAGACGCCAUCCACGCUGUUUUGACCUCCAUAGAAGACACCGGGACCGAUCCAGCCUCCGCGGCCGGGAACGGUGCAUUGGAACGCGGAUUCCCCGUGCCAAGAGUGACUCACCGUCCUUGACACGACACGAUGAUAAUAUGGUGAGCAAGGGCGAGGAGGUCAUCAAAGAGUUCAUGCGCUUCAAGGUGCGCAUGGAGGGCUCCAUGAACGGCCACGAGUUCGAGAUCGAGGGCGAGGGCGAGGGCCGCCCCUACGAGGGCACCCAGACCGCCAAGCUGAAGGUGACCAAGGGCGGCCCCCUGCCCUUCGCCUGGGACAUCCUGUCCCCCCAGUUCAUGUACGGCUCCAAGGCGUACGUGAAGCACCCCGCCGACAUCCCCGAUUACAAGAAGCUGUCCUUCCCCGAGGGCUUCAAGUGGGAGCGCGUGAUGAACUUCGAGGACGGCGGUCUGGUGACCGUGACCCAGGACUCCUCCCUGCAGGACGGCACGCUGAUCUACAAGGUGAAGAUGCGCGGCACCAACUUCCCCCCCGACGGCCCCGUAAUGCAGAAGAAGACCAUGGGCUGGGAGGCCUCCACCGAGCGCCUGUACCCCCGCGACGGCGUGCUGAAGGGCGAGAUCCACCAGGCCCUGAAGCUGAAGGACGGCGGCCACUACCUGGUGGAGUUCAAGACCAUCUACAUGGCCAAGAAGCCCGUGCAACUGCCCGGCUACUACUACGUGGACACCAAGCUGGACAUCACCUCCCACAACGAGGACUACACCAUCGUGGAACAGUACGAGCGCUCCGAGGGCCGCCACCACCUGUUCCUGGGGCAUGGCACCGGCAGCACCGGCAGCGGCAGCUCCGGCACCGCCUCCUCCGAGGACAACAACAUGGCCGUCAUCAAAGAGUUCAUGCGCUUCAAGGUGCGCAUGGAGGGCUCCAUGAACGGCCACGAGUUCGAGAUCGAGGGCGAGGGCGAGGGCCGCCCCUACGAGGGCACCCAGACCGCCAAGCUGAAGGUGACCAAGGGCGGCCCCCUGCCCUUCGCCUGGGACAUCCUGUCCCCCCAGUUCAUGUACGGCUCCAAGGCGUACGUGAAGCACCCCGCCGACAUCCCCGAUUACAAGAAGCUGUCCUUCCCCGAGGGCUUCAAGUGGGAGCGCGUGAUGAACUUCGAGGACGGCGGUCUGGUGACCGUGACCCAGGACUCCUCCCUGCAGGACGGCACGCUGAUCUACAAGGUGAAGAUGCGCGGCACCAACUUCCCCCCCGACGGCCCCGUAAUGCAGAAGAAGACCAUGGGCUGGGAGGCCUCCACCGAGCGCCUGUACCCCCGCGACGGCGUGCUGAAGGGCGAGAUCCACCAGGCCCUGAAGCUGAAGGACGGCGGCCACUACCUGGUGGAGUUCAAGACCAUCUACAUGGCCAAGAAGCCCGUGCAACUGCCCGGCUACUACUACGUGGACACCAAGCUGGACAUCACCUCCCACAACGAGGACUACACCAUCGUGGAACAGUACGAGCGCUCCGAGGGCCGCCACCACCUGUUCCUGUACGGCAUGGACGAGCUGUACAAGUGAACGCGUCUGGAACAAUCGGGUGGCAUCCCUGUGACCCCUCCCCAGUGCCUCUCCUGGCCCUGGAAGUUGCCACUCCAGUGCCCACCAGCCUUGUCCUAAUAAAAUUAAGUUGCAUCAAGCUCUA

copGFP RNA sequence (GenBank: KX757255.1)

GGGAGAGCCGCCACCAUGGAGAGCGACGAGAGCGGCCUGCCCGCCAUGGAGAUCGAGUGCCGCAUCACCGGCACCCUGAACGGCGUGGAGUUCGAGCUGGUGGGCGGCGGAGAGGGCACCCCCAAGCAGGGCCGCAUGACCAACAAGAUGAAGAGCACCAAAGGCGCCCUGACCUUCAGCCCCUACCUGCUGAGCCACGUGAUGGGCUACGGCUUCUACCACUUCGGCACCUACCCCAGCGGCUACGAGAACCCCUUCCUGCACGCCAUCAACAACGGCGGCUACACCAACACCCGCAUCGAGAAGUACGAGGACGGCGGCGUGCUGCACGUGAGCUUCAGCUACCGCUACGAGGCCGGCCGCGUGAUCGGCGACUUCAAGGUGGUGGGCACCGGCUUCCCCGAGGACAGCGUGAUCUUCACCGACAAGAUCAUCCGCAGCAACGCCACCGUGGAGCACCUGCACCCCAUGGGCGAUAACGUGCUGGUGGGCAGCUUCGCCCGCACCUUCAGCCUGCGCGACGGCGGCUACUACAGCUUCGUGGUGGACAGCCACAUGCACUUCAAGAGCGCCAUCCACCCCAGCAUCCUGCAGAACGGGGGCCCCAUGUUCGCCUUCCGCCGCGUGGAGGAGCUGCACAGCAACACCGAGCUGGGCAUCGUGGAGUACCAGCACGCCUUCAAGACCCCCAUCGCCUUCGCCAGAUCCCGCGCUCAGUCGUCCAAUUCUGCCGUGGACGGCACCGCCGGACCCGGCUCCACCGGAUCUCGCUCCCCCAAGAAGAAGAGGAAAGUCUCGAGCGACUACAAAGACCUAUAGGUGGCCCAAUUAAUGACGGUGAUUAUAAAGAUCAUGACAUCGAUUACAAGGAUGACGAUGACAAGGCGCAGGAUGACCGGUCAUCAUCACCAUCACCAUUGAGU

cas9 RNA sequence (Addgene: 72247)

GGGAGAGCCGCCACCAUGGAUAAAAAGUAUUCUAUUGGUUUAGACAUCGGCACUAAUUCCGUUGGAUGGGCUGUCAUAACCGAUGAAUACAAAGUACCUUCAAAGAAAUUUAAGGUGUUGGGGAACACAGACCGUCAUUCGAUUAAAAAGAAUCUUAUCGGUGCCCUCCUAUUCGAUAGUGGCGAAACGGCAGAGGCGACUCGCCUGAAACGAACCGCUCGGAGAAGGUAUACACGUCGCAAGAACCGAAUAUGUUACUUACAAGAAAUUUUUAGCAAUGAGAUGGCCAAAGUUGACGAUUCUUUCUUUCACCGUUUGGAAGAGUCCUUCCUUGUCGAAGAGGACAAGAAACAUGAACGGCACCCCAUCUUUGGAAACAUAGUAGAUGAGGUGGCAUAUCAUGAAAAGUACCCAACGAUUUAUCACCUCAGAAAAAAGCUAGUUGACUCAACUGAUAAAGCGGACCUGAGGUUAAUCUACUUGGCUCUUGCCCAUAUGAUAAAGUUCCGUGGGCACUUUCUCAUUGAGGGUGAUCUAAAUCCGGACAACUCGGAUGUCGACAAACUGUUCAUCCAGUUAGUACAAACCUAUAAUCAGUUGUUUGAAGAGAACCCUAUAAAUGCAAGUGGCGUGGAUGCGAAGGCUAUUCUUAGCGCCCGCCUCUCUAAAUCCCGACGGCUAGAAAACCUGAUCGCACAAUUACCCGGAGAGAAGAAAAAUGGGUUGUUCGGUAACCUUAUAGCGCUCUCACUAGGCCUGACACCAAAUUUUAAGUCGAACUUCGACUUAGCUGAAGAUGCCAAAUUGCAGCUUAGUAAGGACACGUACGAUGACGAUCUCGACAAUCUACUGGCACAAAUUGGAGAUCAGUAUGCGGACUUAUUUUUGGCUGCCAAAAACCUUAGCGAUGCAAUCCUCCUAUCUGACAUACUGAGAGUUAAUACUGAGAUUACCAAGGCGCCGUUAUCCGCUUCAAUGAUCAAAAGGUACGAUGAACAUCACCAAGACUUGACACUUCUCAAGGCCCUAGUCCGUCAGCAACUGCCUGAGAAAUAUAAGGAAAUAUUCUUUGAUCAGUCGAAAAACGGGUACGCAGGUUAUAUUGACGGCGGAGCGAGUCAAGAGGAAUUCUACAAGUUUAUCAAACCCAUAUUAGAGAAGAUGGAUGGGACGGAAGAGUUGCUUGUAAAACUCAAUCGCGAAGAUCUACUGCGAAAGCAGCGGACUUUCGACAACGGUAGCAUUCCACAUCAAAUCCACUUAGGCGAAUUGCAUGCUAUACUUAGAAGGCAGGAGGAUUUUUAUCCGUUCCUCAAAGACAAUCGUGAAAAGAUUGAGAAAAUCCUAACCUUUCGCAUACCUUACUAUGUGGGACCCCUGGCCCGAGGGAACUCUCGGUUCGCAUGGAUGACAAGAAAGUCCGAAGAAACGAUUACUCCCUGGAAUUUUGAGGAAGUUGUCGAUAAAGGUGCGUCAGCUCAAUCGUUCAUCGAGAGGAUGACCGCCUUUGACAAGAAUUUACCGAACGAAAAAGUAUUGCCUAAGCACAGUUUACUUUACGAGUAUUUCACAGUGUACAAUGAACUCACGAAAGUUAAGUAUGUCACUGAGGGCAUGCGUAAACCCGCCUUUCUAAGCGGAGAACAGAAGAAAGCAAUAGUAGAUCUGUUAUUCAAGACCAACCGCAAAGUGACAGUUAAGCAAUUGAAAGAGGACUACUUUAAGAAAAUUGAAUGCUUCGAUUCUGUCGAGAUCUCCGGGGUAGAAGAUCGAUUUAAUGCGUCACUUGGUACGUAUCAUGACCUCCUAAAGAUAAUUAAAGAUAAGGACUUCCUGGAUAACGAAGAGAAUGAAGAUAUCUUAGAAGAUAUAGUGUUGACUCUUACCCUCUUUGAAGAUCGGGAAAUGAUUGAGGAAAGACUAAAAACAUACGCUCACCUGUUCGACGAUAAGGUUAUGAAACAGUUAAAGAGGCGUCGCUAUACGGGCUGGGGAGCCUUGUCGCGGAAACUUAUCAACGGGAUAAGAGACAAGCAAAGUGGUAAAACUAUUCUCGAUUUUCUAAAGAGCGACGGCUUCGCCAAUAGGAACUUUAUGGCCCUGAUCCAUGAUGACUCUUUAACCUUCAAAGAGGAUAUACAAAAGGCACAGGUUUCCGGACAAGGGGACUCAUUGCACGAACAUAUUGCGAAUCUUGCUGGUUCGCCAGCCAUCAAAAAGGGCAUACUCCAGACAGUCAAAGUAGUGGAUGAGCUAGUUAAGGUCAUGGGACGUCACAAACCGGAAAACAUUGUAAUCGAGAUGGCACGCGAAAAUCAAACGACUCAGAAGGGGCAAAAAAACAGUCGAGAGCGGAUGAAGAGAAUAGAAGAGGGUAUUAAAGAACUGGGCAGCCAGAUCUUAAAGGAGCAUCCUGUGGAAAAUACCCAAUUGCAGAACGAGAAACUUUACCUCUAUUACCUACAAAAUGGAAGGGACAUGUAUGUUGAUCAGGAACUGGACAUAAACCGUUUAUCUGAUUACGACGUCGAUCACAUUGUACCCCAAUCCUUUUUGAAGGACGAUUCAAUCGACAAUAAAGUGCUUACACGCUCGGAUAAGAACCGAGGGAAAAGUGACAAUGUUCCAAGCGAGGAAGUCGUAAAGAAAAUGAAGAACUAUUGGCGGCAGCUCCUAAAUGCGAAACUGAUAACGCAAAGAAAGUUCGAUAACUUAACUAAAGCUGAGAGGGGUGGCUUGUCUGAACUUGACAAGGCCGGAUUUAUUAAACGUCAGCUCGUGGAAACCCGCGCCAUCACAAAGCAUGUUGCGCAGAUACUAGAUUCCCGAAUGAAUACGAAAUACGACGAGAACGAUAAGCUGAUUCGGGAAGUCAAAGUAAUCACUUUAAAGUCAAAAUUGGUGUCGGACUUCAGAAAGGAUUUUCAAUUCUAUAAAGUUAGGGAGAUAAAUAACUACCACCAUGCGCACGACGCUUAUCUUAAUGCCGUCGUAGGGACCGCACUCAUUAAGAAAUACCCGAAGCUAGAAAGUGAGUUUGUGUAUGGUGAUUACAAAGUUUAUGACGUCCGUAAGAUGAUCGCGAAAAGCGAACAGGAGAUAGGCAAGGCUACAGCCAAAUACUUCUUUUAUUCUAACAUUAUGAAUUUCUUUAAGACGGAAAUCACUCUGGCAAACGGAGAGAUACGCAAACGACCUUUAAUUGAAACCAAUGGGGAGACAGGUGAAAUCGUAUGGGAUAAGGGCCGGGACUUCGCGACGGUGAGAAAAGUUUUGUCCAUGCCCCAAGUCAACAUAGUAAAGAAAACUGAGGUGCAGACCGGAGGGUUUUCAAAGGAAUCGAUUCUUCCAAAAAGGAAUAGUGAUAAGCUCAUCGCUCGUAAAAAGGACUGGGACCCGAAAAAGUACGGUGGCUUCGAUAGCCCUACAGUUGCCUAUUCUGUCCUAGUAGUGGCAAAAGUUGAGAAGGGAAAAUCCAAGAAACUGAAGUCAGUCAAAGAAUUAUUGGGGAUAACGAUUAUGGAGCGCUCGUCUUUUGAAAAGAACCCCAUCGACUUCCUUGAGGCGAAAGGUUACAAGGAAGUAAAAAAGGAUCUCAUAAUUAAACUACCAAAGUAUAGUCUGUUUGAGUUAGAAAAUGGCCGAAAACGGAUGUUGGCUAGCGCCGGAGAGCUUCAAAAGGGGAACGAACUCGCACUACCGUCUAAAUACGUGAAUUUCCUGUAUUUAGCGUCCCAUUACGAGAAGUUGAAAGGUUCACCUGAAGAUAACGAACAGAAGCAACUUUUUGUUGAGCAGCACAAACAUUAUCUCGACGAAAUCAUAGAGCAAAUUUCGGAAUUCAGUAAGAGAGUCAUCCUAGCUGAUGCCAAUCUGGACAAAGUAUUAAGCGCAUACAACAAGCACAGGGAUAAACCCAUACGUGAGCAGGCGGAAAAUAUUAUCCAUUUGUUUACUCUUACCAACCUCGGCGCUCCAGCCGCAUUCAAGUAUUUUGACACAACGAUAGAUCGCAAACGAUACACUUCUACCAAGGAGGUGCUAGACGCGACACUGAUUCACCAAUCCAUCACGGGAUUAUAUGAAACUCGGAUAGAUUUGUCACAGCUUGGGGGUGACGGAUCCCCCAAGAAGAAGAGGAAAGUCUCGAGCGACUACAAAGACCAUGACGGUGAUUAUAAAGAUCAUGACAUCGAUUACAAGGAUGACGAUGACAAGGCUGCAGGAUGACCGGUCAUCAUCACCAUCACCAUUGAGU

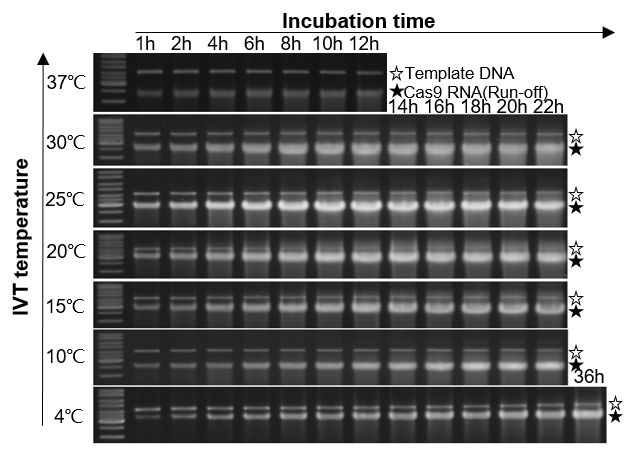

**Figure S1.** Yield of cas9 RNA in VSW-3 RNAP IVT with various incubation temperature and time. The gel bands corresponding to DNA templates were indicated by empty stars and those corresponding to run-off cas9 RNA indicated by filled stars. Maximum yield was obtained at 25℃ for 12 hours. At lower temperatures, extended incubation increased the yield.

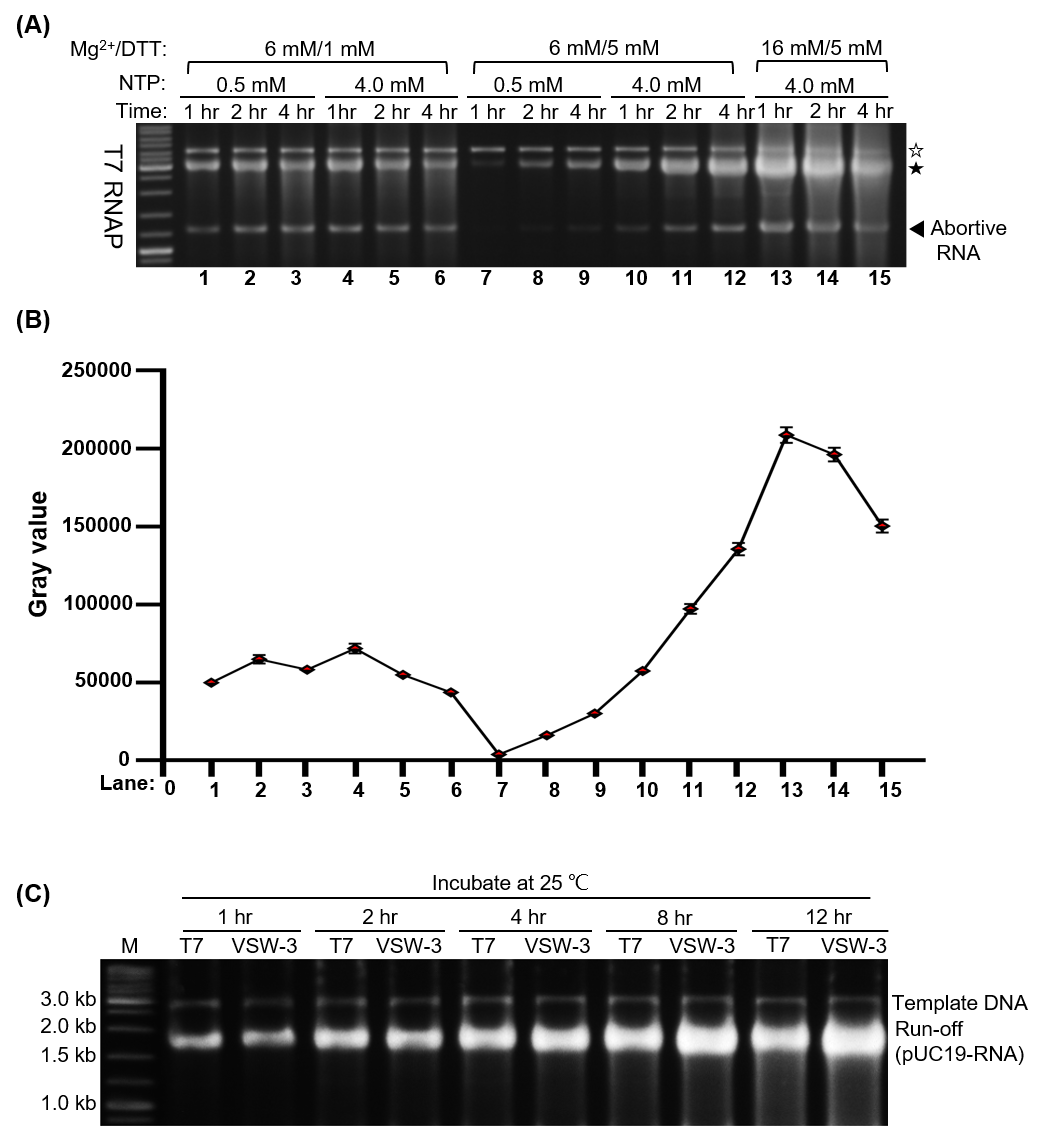

**Figure S2.** Optimal IVT conditions for T7 RNAP. **(A)** The optimized MgCl_2_/NTP/DTT concentration (16/4/5 mM) for VSW-3 RNAP also applies to T7 RNAP. While extended incubation time (>1 hour) decreases the yield of run-off transcripts of T7 RNAP at 37℃. The gel bands corresponding to DNA templates were indicated by empty star and those corresponding to run-off cas9 RNA indicated by filled star. **(B)** Gray-scale quantitation of the run-off cas9 RNA in **(A)** by image J software revealed that T7 RNAP reaches its maximum yield in 1 hour with the optimized IVT conditions at 37℃. **(C)** Comparison of the IVT yield (pUC19-RNA) between T7 RNAP and VSW-3 RNAP at 25℃, with the same IVT conditions and at the same enzyme and substrates concentration.

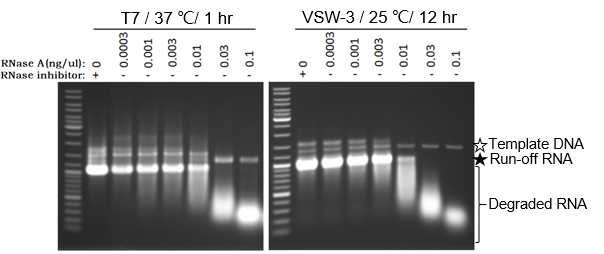

**Figure S3.** RNA degradation in the presence of RNase A in T7 and VSW-3 RNAP IVT. With the same level of added RNase A, RNA transcript degradation is more severe in T7 RNAP IVT (37℃, 1 hour) than that in VSW-3 RNAP IVT (25℃, 12 hours).

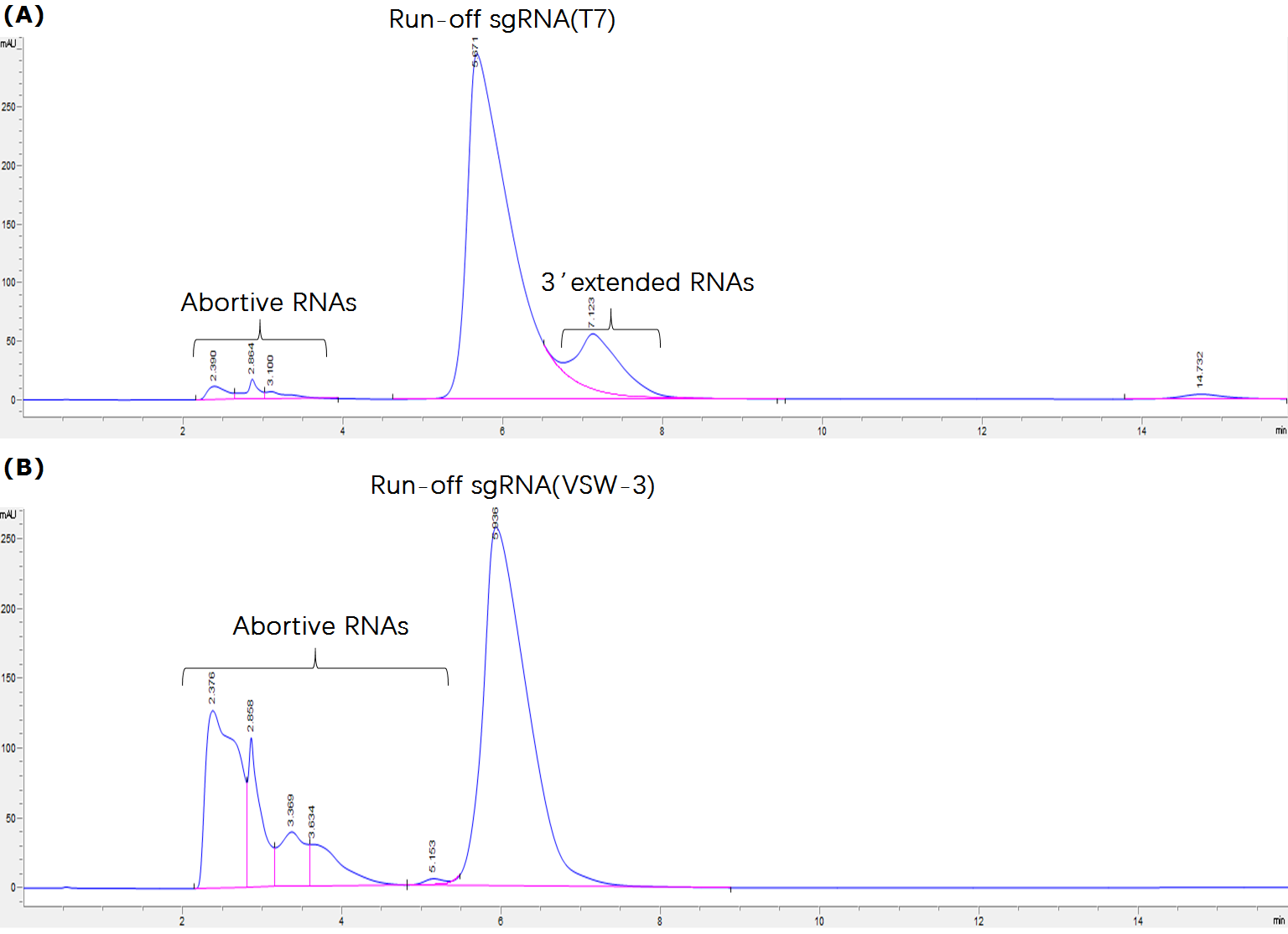

**Figure S4.** HPLC analysis of the sgRNA transcripts from T7 (A) and VSW-3 (B) RNAP IVT. **(A)** The HPLC chromatogram of T7 sgRNA transcripts contains a small peak (corresponding to 3’-extended transcripts) following the main peak (corresponding to run-off transcripts). **(B)** The HPLC chromatogram of VSW-3 sgRNA transcripts shows no peak following the main peak, supporting the conclusion from the PAGE and 3’ RACE that VSW-3 RNAP does not produce 3’-extended transcripts. However, peaks corresponding to abortive products are more significant in VSW-3 chromatogram.
